# Towards a Cytometry Foundation Model: Interpretable Sample-level Predictive Modelling via Pretrained Transformers

**DOI:** 10.64898/2026.03.31.712806

**Authors:** Zixin Zhuang, Benjamin S. Mashford, Liang Zheng, T. Daniel Andrews

## Abstract

Foundation models have transformed scientific data modelling across domains, yet flow cytometry has lacked one. Despite the abundance of high-dimensional cellular data, automated analysis remains bottlenecked by marker variability: prior studies are typically confined to fixed marker panels and homogeneous data, limiting scalability and generalisation due to architectural constraints. We present the Generalised Pretrained Cytometry Transformer (GPCT), an interpretable framework designed to learn from heterogeneous marker panels for sample-level predictive modelling. Through a novel cytometry-specific pretraining regime, GPCT learns transferable cellular representations that achieve high classification accuracy across diverse datasets. Notably, pretraining significantly boosts performance on data-scarce downstream tasks, marking a pivotal step towards a cytometry foundation model. Furthermore, GPCT maintains interpretability and identifies the specific cell subsets most influential to its predictions. This enables direct biological validation of learned patterns and provides a data-driven basis for refining traditional gating strategies.

## 1 Introduction

Flow cytometry is an affordable, accessible method for ascertaining the cellular phenotype of an organism, often from just a small peripheral blood samples. Each cell passing through the cytometer is represented by a signature of marker intensity values across the panel, quantifying protein (markers) expression at single-cell resolution. Historically, analysis has been manual and relied on visual inspection of pairwise marker intensity plots — a process with a combinatorial complexity now exceeds human capacity as modern flow and spectral flow cytometers push panel sizes ever larger.

In standard analysis, cell types are identified through manual gating: experts draw polygons around cell clusters on a aforementioned 2D scatter plot of marker intensity, then repeat this process across successive marker pairs, progressively filtering down to isolate specific cell populations. This becomes increasingly time-consuming and potentially more biased as panel sizes grow. Furthermore, strong batch effects across runs, machines, and sites prevent universal gate sets, requiring manual recalibration for each new context. Substantial variation in marker antibodies between experiments — driven largely by experimentalist preference — also results in inconsistent gates and signals, often forcing channels to be discarded.

A computational alternative to manual gating is to automatically capture cell types and patterns using Machine Learning (ML). Unsupervised learning algorithms [1] require no external labels, and are commonly used to identify distinct cell types and build automatic gates, though they typically suffer from the same data inconsistency issues described above [2]. Supervised algorithms, on the other hand, correlate flow data with external labels — such as disease phenotypes or genotypes. By modelling the relationship between cellular phenotype and sample-level labels, they enable end-to-end analysis of patient conditions.

Notably, deep-learning-based methods across a range of architectures (CNN [3, 4], Transformer encoder [5, 6], MLP with pooling [7–9], and GNN [10]) can directly process raw marker expression values through learned representation, negating the need for difficult, lossy and bias-prone manual feature extraction. However, data and marker inconsistency remain persistent challenges, alongside direct sample-level prediction and inherent model interpretability [2]. Together, these factors have confined most ML-based methods to datasets with consistent marker panels, which are often small in size. Prior studies have also tended to focus on homogeneous datasets with landmark cellular phenotypes, such as those observed in leukaemia [5, 8]. Meanwhile, detection of subtle changes in minor cell populations remains crucial for clinical applications and a challenge future methods must address.

The transformer architecture [11] has been the backbone of Large Language Models (LLMs) and contemporary Artificial Intelligence [12–14]. The attention-mechanism-backed architecture learns patterns from set-structured inputs, making it suitable for a range of data types across scientific and biological domains. Modern LLMs typically incorporates a pretraining stage [12], where the model learns robust representations from unlabelled input data via self-supervision. The resulting model, often termed a foundation models, can be used for improving performance on downstream tasks, especially when downstream task-specifc data is scarce.

In light of the challenges facing ML in cytometry [2], we draw inspiration from the success of LLMs [15] and propose the Generalised Pretrained Cytometry Transformer (GPCT): an end-to-end sample-level predictive modelling pipeline for flow cytometry. GPCT comprises of an interpretable encoder-decoder transformer and a simple yet effective embedding that learns fixed-size features from variable marker panels (Fig. 1).

**Fig. 1.**
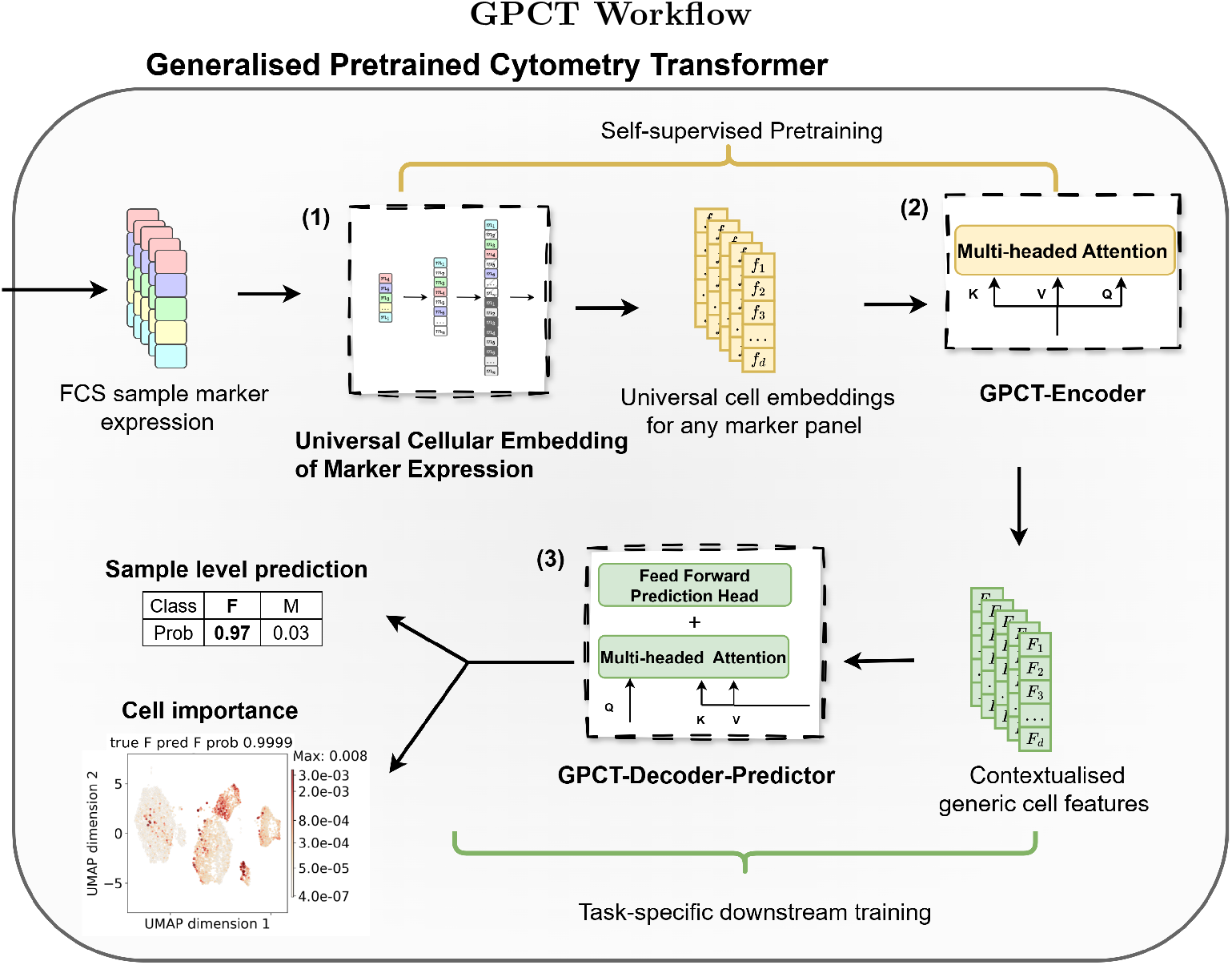
The workflow of GPCT demonstrated on a binary classification task: (1) Sample marker expressions are extracted from FCS files and projected into a fixed-size universal cell representation space applicable to all marker panels (2) GPCT-Encoder extracts implicit cellular patterns and contextualises cellular features through self-attention (3) GPCT-Decoder-Predictor uses the crossattention between task-specific information and learned generic cell features to make a sample level prediction. The relevance of each cell with respect to model prediction is explainable through attention weights.

GPCT has three primary features:

1. Cross-panel compatibility: It natively accommodates inconsistent marker panels across heterogenous data, without requiring separate models for each.
2. Scalable self-supervised pretraining: It employs a cytometry-specific pretraining scheme that learns from vast quantities of data without consistent labelling, capturing robust cellular patterns that boosts performance on downstream tasks.
3. Cell-level interpretability of sample-level predictions: The attention mechanism incorporated in GPCT enables direct sample-level predictions from single-cell inputs, while attributing prediction relevance back to each individual cell.

We demonstrate GPCT on two independent flow cytometry datasets from mutant laboratory mice. Together, they reflect challenges common to datasets in the field: inconsistent marker usage and methodological variation over time for integrated datasets, and data scarcity for small datasets. Across the two datasets, GPCT consistently predicts biological sex or the genetic alteration present in mouse pedigrees. Finally, leveraging the biological sex label that is common to both, we investigate out-of-distribution generalisation on these two vastly different datasets, exploring whether knowledge learned from a larger dataset can be reliably transferred to benefit performance on a smaller dataset.

## 2 Results

### 2.1 GPCT architecture enables modelling across heterogenous marker panels

GPCT (Fig. 1) handles heterogenous marker panels through the Universal Cellular Embedding of Marker expression (UCEM embedding), a novel learnable embedding designed to unambiguously capture per-cell marker expression from any marker panel, and return a fixed-size cellular representation. The encoded cells are then passed on to a modified Transformer encoder-decoder architecture (See Methods for details), where self-attention contextualises the UCEM embedding to extract biologically meaningful features more robust to information misalignment due to marker inconsistency. These features are ultimately used for making sample-level predictions.

The GPCT pipeline is end-to-end differentiable and is trained in two stages: 1. the self-supervised pretraining stage that optimises cellular representation learning, and 2. the subsequent task-specific downstream training that focuses on inferring the correct sample-level labels.

In pretraining, GPCT optimises the UCEM embedding and the GPCT-Encoder via masked prediction on unlabelled data. By learning complex marker co-expression patterns in an uncertainty-aware manner, GPCT promotes comparability of different marker panels by constructing a shared latent representation space. During downstream training, this latent space is preserved via freezing the UCEM embedding and the GPCT-Encoder. Conversely, the GPCT-Decoder-Predictor is trained to infer sample level labels based on the encoder output by minimising task-specific loss. Henceforth, the decoder-predictor will be simply referred to as the “decoder”. During inference, the decoder generates per-cell attention weights indicative of relative cell importance to the predictions, enabling the identification of cell populations relevant to a specific task and its associated biological label(s).

Here, we adopt the term “decoder” following prior work on set-structured data modelling [16] and cytometry data modelling [17], to refer to the stack of crossattention blocks that extracts task-relevant discriminative signals from latent cellular representations, rather than in a generative or reconstructive sense. For convenience, we extend this term to include the attached prediction head. the decoder-predictor module may equivalently be referred to as an “iterative cross-attention pooling based discriminator”.

### 2.2 GPCT models diverse flow cytometry immunophenotyping datasets

#### Dataset 1: Longitudinal mouse immunophenotype dataset

As part of a long-running mutagenesis project to investigate novel genetic causes of immune dysfunction [18], flow cytometry phenotypes for over forty thousand C57BL/6 mice were obtained at the Australian Phenomics Facility between 1995 and 2015. This data is comprised of predominantly eight-colour experiments with varying marker/antibody/fluorophore combinations, yet most samples include a backbone of six common markers (IgM, IgD, B220, CD44, CD4, CD3) (Supplementary Table 10 and 8).

In the present analysis, we have chosen a subset of 14,014 flow cytometry samples (6,978 female, 7,036 male) with a consistent gender metadata label and mostly pan-leukocyte marker panels. Sexual dimorphism rarely produces landmark cell populations readily detectable by manual analysis of flow cytometry data. However, this has proven a tractable problem with application of neural networks [19], with discriminative signals usually subtle and dispersed across multiple cell populations.

#### Dataset 2: Knockout Mouse Project immunophenotype dataset

The Knockout Mouse Project (KOMP) [20] generated mouse strains harbouring gene knockouts for the majority of genes in the mouse genome, accompanied by phenotype data including flow cytometry information for a subset of mutant mouse lines. For our purposes, we focus on a subset of samples subjected to flow cytometry assay of a T cell immunophenotyping panel [21] (Supplementary Table 3). Despite containing nearly 7000 samples, this dataset poses a classic lack-of-data problem, as each knockout (KO) is represented by only 10 to 20 samples. As most knockouts in this dataset were found to lack discernible cellular phenotypes [21], we selected just 5 knock-out lines with clear mutant phenotypes characterised by the original study. This yields 72 samples (Supplementary Table 9) for a 5-class KO classification task.

#### Modelling biological sex-associated cellular patterns across Datasets 1 and 2

A particularly interesting scenario arises in building a single predictive model for both datasets, which share 5 common markers (Supplementary Table 2) and both have sample-level labels indicating biological sex. Specifically, we aim to simulate a smalldata setting with Dataset 2 by reserving most of its samples for testing, and only a small subset for training (See supplementary information for details). We will be exploring the effects of training with a mixture of data from both datasets across the two training stages of GPCT in the following two scenarios:

1. The generalisation scenario, where the model has no access to Dataset 2 labels, is primarily trained with Dataset 1, and evaluated on Dataset 2.
2. Learning with small data scenario, where the model has access to the small subset of labelled Dataset 2 and the labelled Dataset 1, and evaluated on the remaining Dataset 2 samples.

### 2.3 GPCT accurately classifies biological sex across heterogeneous input data

In this section, we evaluate GPCT on biological sex classification using both datasets. We first assess the role of encoder pretraining to classification performance and cross-panel robustness using Dataset 1, then examine the interpretability of GPCT predictions through attention maps, before finally investigating cross-dataset modelling and knowledge transfer between Dataset 1 and 2.

#### 2.3.1 Pretraining drives performance gains and cross-panel robustness on Dataset 1

Technically, the pretraining stage and the GPCT-encoder are not strictly required for the GPCT pipeline to function as intended and produce interpretable sample-level predictions from raw cytometry data. The encoder and the UCEM embedding can be trained jointly with the decoder (*Encoder not Pretrained*), or omitted entirely (*Decoder Only*), with GPCT-Decoder taking the cell embeddings directly as input. However, as evident in Table 1, GPCT with encoder pretraining (*Encoder Pretrained*) achieves superior 7-fold-cross-validated performance under the same training conditions, outperforming both alternatives and achieving 87% accuracy and an Area Under the Curve (AUC) score of 0.938 across all samples. Notably, incorporating a non-pretrained encoder is shown to decrease the performance relative to the decoder-only model, likely due to the increased model complexity that hinders optimisation. ROC curves and confusion matrices for all three models can be found in Supplementary Fig. 3.

**Table 1.** The 7-fold batch-grouped cross-validated performance of three GPCT variations: Decoder Only (without encoder), GPCT with non-pretrained encoder, and full GPCT with encoder pretraining.

| Cross-validated GPCT performance for biological sex classification |  |  |  |  |
| --- | --- | --- | --- | --- |
| Model | Uses encoder | Encoder pretrained | ACC | AUC |
| Decoder Only | No | No | 0.838 | 0.906 |
| Encoder not Pretrained | Yes | No | 0.802 | 0.876 |
| Encoder Pretrained | Yes | Yes | **0.870** | **0.938** |

We then verify that GPCT can generalise not only to unseen samples, but also to unseen marker panels. In a leave-one-panel-out experiment (training mode A in Table 2), we selected the two most represented panels in the dataset to form the test set in two separate runs, and trained GPCT on all remaining panels respectively. In both cases, GPCT achieved less than 8% performance drop compared to the baseline (training mode B), where the respective test panel was seen during both training stages. This demonstrates that GPCT is reasonably robust to moderate variations in marker panels. Conversely, we performed a companion experiment (training mode C in Table 2), training and evaluating only on samples from the respective panel under the same 7-fold cross validation setting as training mode B. Compared with mode C, training mode B utilises the full dataset and achieves a better performance on those panel-specific samples. This further demonstrates that GPCT not only enables modelling across heterogeneous marker panels, but also effectively leverages the additional information learned from different panels to improve panel-specific predictions.

**Table 2.** GPCT performance of samples from two most represented marker panels, under three different training modes: (1) *Leave panel out* sets aside all samples from the panel as the testing set, with the model trained using rest of the samples belonging to different marker panels. (2) *Full dataset cross val* follows the same cross validation procedure as in Table 1, using the full Dataset 1. Predictions for samples from the specified panel are collected and aggregated to calculate the metrics. (3) *Panel samples cross val* follows the same cross validation split as *Full dataset cross val*, but includes only samples from the specified panel in training and testing.

| Marker Panel | Training Mode | ACC | AUC | Panel samples |
| --- | --- | --- | --- | --- |
| B220, CD3, CD4, CD44, IgD, IgM, KLRG1, NK1-1 | A. Leave panel out | 0.798 | 0.878 | 7002 |
|  | B. Full dataset cross val | 0.865 | 0.938 |  |
|  | C. Panel samples cross val | 0.851 | 0.920 |  |
| B220, CD25, CD3, CD4, CD44, IgD, IgM, NK1-1 | A. Leave panel out | 0.825 | 0.904 | 6013 |
|  | B. Full dataset cross val | 0.893 | 0.953 |  |
|  | C. Panel samples cross val | 0.870 | 0.934 |  |

#### 2.3.2 Attention weights identify biologically relevant cell populations

Interpretability is essential for validating the biological basis of model predictions and discovering the specific cellular subsets that drive sample-level outcomes. In this regard, GPCT can be interpreted through the attention mechanism used by the decoder: during inference, each attention head in the multi-head attention layer assigns a weight to every cell, representing its relative contribution to the decision-making process. These weights serve as a quantitative measure of per-cell “importance”, and while they are typically averaged across heads per layer for visualisation, each layer may capture distinct patterns that reflect the model’s internal processing steps.

For the biological sex prediction, we observed distinct cellular attention patterns amongst the four attention layers in the GPCT-Decoder (Fig. 2a, layers numbered using 0-indexing). Cells are visualised on a 2D plane using UMAP[22] according to their marker expression values. In general, attention tends to be sparse, spanning multiple magnitudes in terms of weights assigned to each cell, except for the second layer (layer 1), where attentions appear to be much more uniform. Cells with high importance often cluster together in the reduced dimensions, sometimes as localised groupings in a larger cell population (e.g. the left most UMAP grouping in the blue circle, identified as IgM^+^ IgD^+^ B cells). Notably, the cell population in the green circle at the bottom right corner maintains high relevance in all but the second layer, and is identified to be NK1-1^+^ KLRG1^+^ cells. The small cell population in the middle purple circle also receives high attention from the three layers, and shows high or elevated expression across a range of markers, including B220, CD3, CD4, CD44, IgM, KLRG1, and NK1-1.

**Fig. 2.**
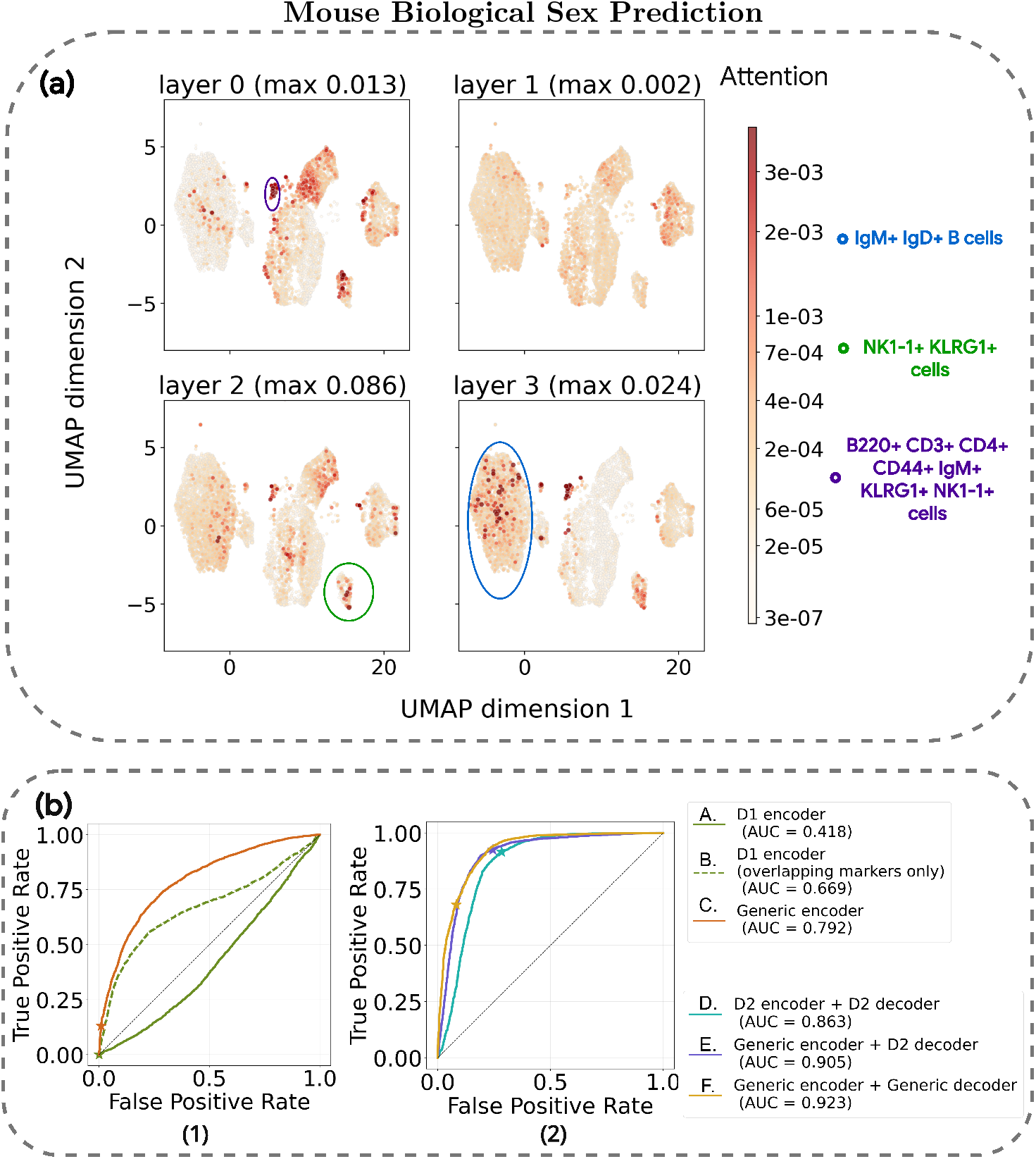
(a) GPCT attention per decoder layer for a correctly classified female sample, visualised using UMAP. Cells are coloured by attention weights, which are averaged over 4 attention heads and sum up to one per layer, with the top and bottom 0.2% attention values clipped for visual colour mapping on a power scale. The largest attention value for each layer is as indicated. There are 7000 cells in total, making the expectation of attention per cell approximately 1.4 ×10^*−*4^. (b) The ROC curves depicting (1) D1-GPCT models generalising to Dataset 2 without prior knowledge of labelled samples from Dataset 2 (2) Performance evolution of D2-GPCT models when samples from Dataset 1 are gradually included in training. The star marker on each curve denotes the classification threshold of 0.5.

In general, highly relevant cells are present across most major cell populations, indicating a complex and broad-spectrum immune signature of biological sex captured by the decision process. The attention pattern shown above remains largely consistent across the dataset, with slight per-sample variation(see Supplementary Fig. 4). Notably, attention tends to be more focused in layer 1 for male samples, often highlighting NK1-1^+^ KLRG1^+^ cells compared with female samples.

#### 2.3.3 Cross-dataset modelling facilitates knowledge transfer across heterogeneous cohorts

We evaluated the impact of cross-dataset pretraining on the model generalisation scenario using two configurations. The first model, the D1 encoder (Experiment A and B), was trained exclusively on Dataset 1. The second, the generic encoder (Experiments C), was pretrained on combined training data from Datasets 1 and 2 before downstream training on Dataset 1 only. Results in Fig. 2b (1) demonstrate that including even a small fraction of Dataset 2 in the pretraining phase significantly improved downstream generalisation to Dataset 2 testing samples. This suggests that multidataset pretraining facilitates the learning of a shared cellular representation, which implicitly aligns the disparate datasets. By leveraging this aligned latent space, the model adopts a decision-making process that transfers more effectively to unseen data. Further information on Experiment A and B can be found in the supplementary information.

For the second scenario, we evaluated the effect of incorporating external Dataset 1 samples across both training stages using three configurations. Experiment D was trained solely on Dataset 2 (D2 encoder and D2 decoder), Experiment E incorporated Dataset 1 during pretraining (generic encoder), and Experiment F extended this to downstream training (generic decoder). Results in Fig. 2b (2) show that GPCT can effectively extract transferable knowledge from additional out-of-distribution data in both training stages, in turn improving performance on in-distribution samples. This demonstrates GPCT’s potential as a foundation model through its ability to extract generalisable signals from diverse data.

### 2.4 GPCT identifies different gene knockouts solely from flow cytometry data

Four models, including A. *CellCnn* [3], B. *GPCT decoder only*, C. *GPCT*, and D.*GPCT with generic encoder*, were trained to classify five different gene KOs in Dataset 2. Leave-one-batch-out cross validation (totalling 31 batches) was employed to avoid false confidence in model performance when the model merely learns the batch effects. All models used only assigned training data per split, except for model D, whose encoder was pretrained on 6,904 mice from Dataset 2 that don’t belong to the five KOs of interest. The results are summarised in Fig. 3, with per-class ROC curves and confusion matrices available in Supplementary Fig. 5, and a summary of performance metrics in Supplementary Table 1.

**Fig. 3.**
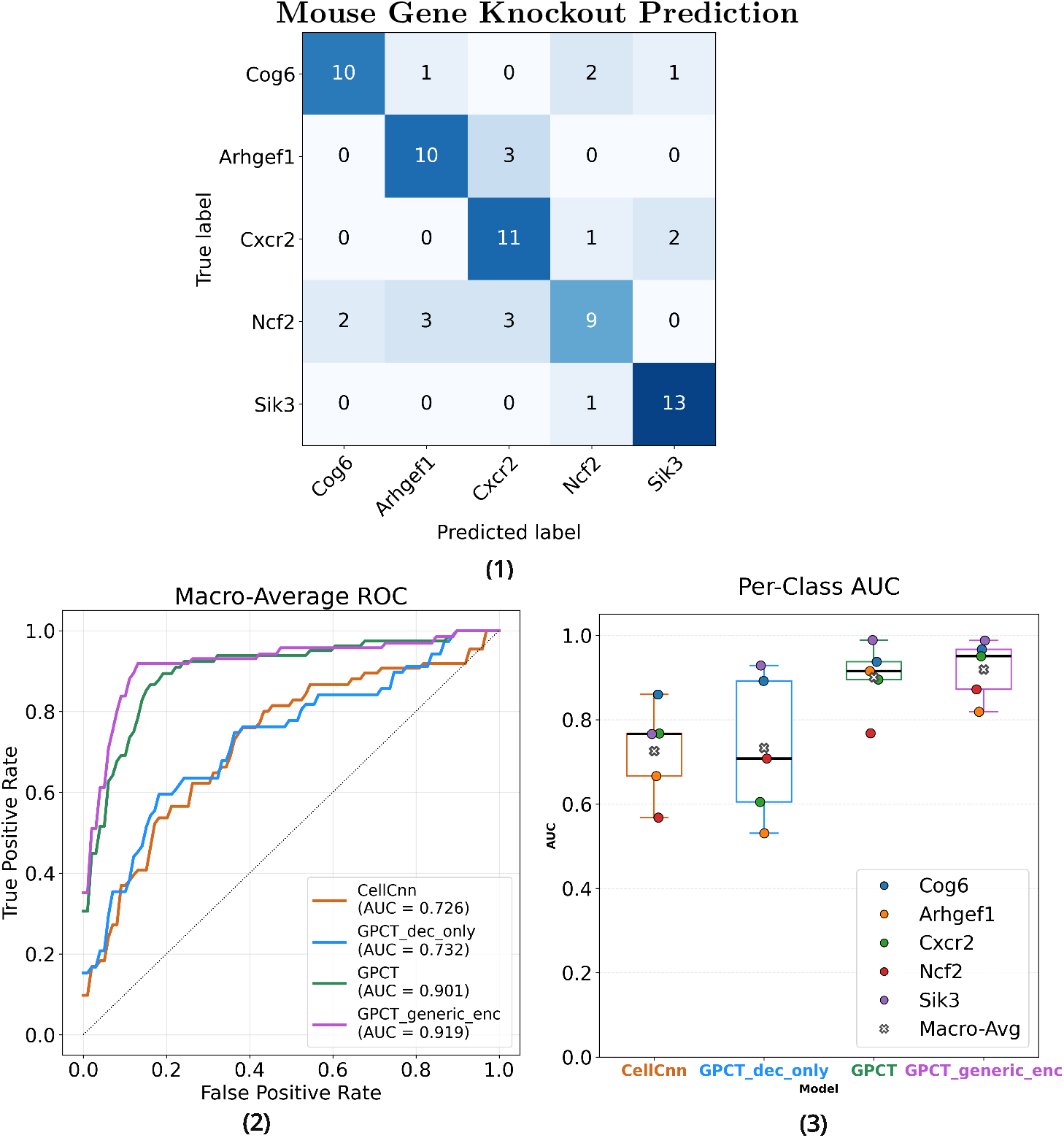
(1) Confusion matrix of the best performing model, GPCT with pretrained generic encoder (2) Macro-average ROC curves of each model, labelled with the one-vs-rest macro averaged AUC score. (3) The distribution of one-vs-rest AUC score per gene KO class for each model

In general, every model outperforms the random classifier baseline across all classes, with Arhgef1 and Ncf2 being the two most difficult KOs to identify, and Sik3 and Cog6 the easiest. Model D, which possesses a generic encoder, is the best performing model overall, achieving a macro-average AUC score of 0.919 and a classification accuracy of 0.736 (confusion matrix at Fig. 3 (1)). model C achieved the best classification performance without any external data, closely behind model D and outperforming models A and B by a significant margin. The decoder-only variant B, lacking an encoder, incorporates no context-aware representation learning, much like CellCnn. Its performance, much closer to that of CellCnn than to the other GPCT models, provides strong evidence that a robust sample representation is crucial for learning with small data. Furthermore, Model D outperforms model C even when the encoder is pretrained without seeing downstream task samples, suggesting that larger-scale pretraining on similarly but not identically distributed data creates a generic and stable representation that rivals task-specific pretraining.

To further investigate the effects of large-scale pretraining, we performed a few-shot analysis on the same task (see Supplementary Figure 6), training GPCT models C and D with Dataset 2 subsets containing different numbers of labelled samples per class (from 1 to 8 per class). Both models improve with more training data, though performance gains are unstable as sample size increased. Model D outperforms model C by a large margin throughout, matching the latter’s best performance even at 1-hot. Crucially, model D consistently performed above random while model C remained close to the random baseline, suggesting that transferred knowledge learned from external data is essential for successful few-shot predictions, even though pretraining itself is not task-discriminative. Nevertheless, both models fall substantially short of their respective leave-one-batch-out performance, highlighting that few-shot cytometry modelling remains a challenge.

## 3 Discussion

These results demonstrate the potential for a cytometry foundation model: via largescale GPCT pretraining. GPCT provides an end-to-end, interpretable pipeline that addresses fundamental challenges in cellular phenotype modelling using cytometry — from marker panel heterogeneity to data scarcity — as demonstrated using the two mouse immunophenotyping datasets. Notably, large-scale pretraining yielded considerable performance gains in small-data settings, attributable to robust cellular representations that recover biological signals despite marker inconsistency. Extending this paradigm to the full corpus of available flow cytometry data could establish a comprehensive repository of generic cellular patterns, transferable to any downstream task — analogous to how foundation models in language and vision benefit data-scarce applications.

Importantly, GPCT also enables the interpretation of predictions through the decoder’s cross-attention weights, which identify the cell populations most relevant to each model decision. This capacity for cell-level attribution can directly inform the refinement of manual gating strategies and facilitate the discovery of rare or novel cell populations associated with specific biological conditions.

Application to the gene knockout prediction task further demonstrates the possibility of correlating genotype with flow cytometry phenotype, which may offer a data-driven basis for characterising disease-associated immune profiles in clinical precision medicine. By natively accommodating marker panel heterogeneity, GPCT enables integration of disparate clinical datasets that were previously difficult to compare, while its automated feature extraction capability reduces the subjectivity inherent to manual gating. Together, these properties position GPCT as a step toward more unified and scalable immune profiling, especially in the realisation of a cytometry foundation model.

Several directions for future work follow naturally from these findings. While GPCT’s pretraining design accounts for experimental batch effects and fluorophore inconsistencies, it does not explicitly reduce these biologically irrelevant sources of variation. Integrating dedicated batch alignment strategies [23] into the pipeline represents a promising extension. Additionally, applying more expressive interpretability methods to the GPCT encoder-decoder [24, 25] could yield more concise and biologically informative cell-level relevance attribution beyond attention weights. Ultimately, applying GPCT to human clinical datasets, where panel diversity and sample scarcity are most acute, represents the natural next step toward establishing it as a practical tool in both diagnostic and research workflows.

## 4 Methods

### 4.1 Problem formulation

For the purpose of this study, each flow cytometry sample *s* is defined as a finite set of cells 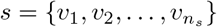, where each cell is featurised by its marker expression vector 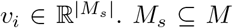. *M*_*s*_ ⊆*M* is the set of cytometry markers measured for sample *s*, and a subset of the finite marker vocabulary *M* containing all possible cytometry markers.

GPCT is framed under a supervised machine learning scenario, where the model aims to learn a mapping *f* : 𝒳 → 𝒴 between each sample *x*_*s*_ ∈ 𝒳 and corresponding sample level label(s) *y*_*s*_ ∈𝒴. The label space 𝒴 can be discrete (classification) or continuous (regression).

Following terminology conventions in deep learning, a vector that represents a cell (can be raw marker expression or learned representation) is also referred to as a *token*, with each element of the token referred to as a *feature*. A cytometry sample containing observations of *n*_*s*_ cells on |*M*_*s*_| markers can be represented as an *n*_*s*_ ×|*M*_*s*_| matrix *X*, though no specific row order is assumed, and any permutations of columns are equivariant.

### 4.2 GPCT model architecture

#### 4.2.1 Universal Cellular Embedding of Marker Expression

The transformer architecture, like most deep learning architectures, requires input tokens to have consistent features, which demands a mechanism to produce fixed size cellular embeddings for any sample marker set *M*_*s*_.

A important characteristic of cytometry data is that not every marker expressed is measured, and unmeasured markers might still exist on the cell surface. To address this issue, we employ a simple yet effective embedding (Fig. 4) that constructs an augmented cell representation by concatenating a one-hot marker availability indicator to a sparse marker expression vector. For each sample *X* ∈ ℝ^*n×m*^ containing *n* cells and *m* measured markers, let *M*_*s*_ denote the set of measured markers and *M* the full marker vocabulary. We define:

**Fig. 4.**
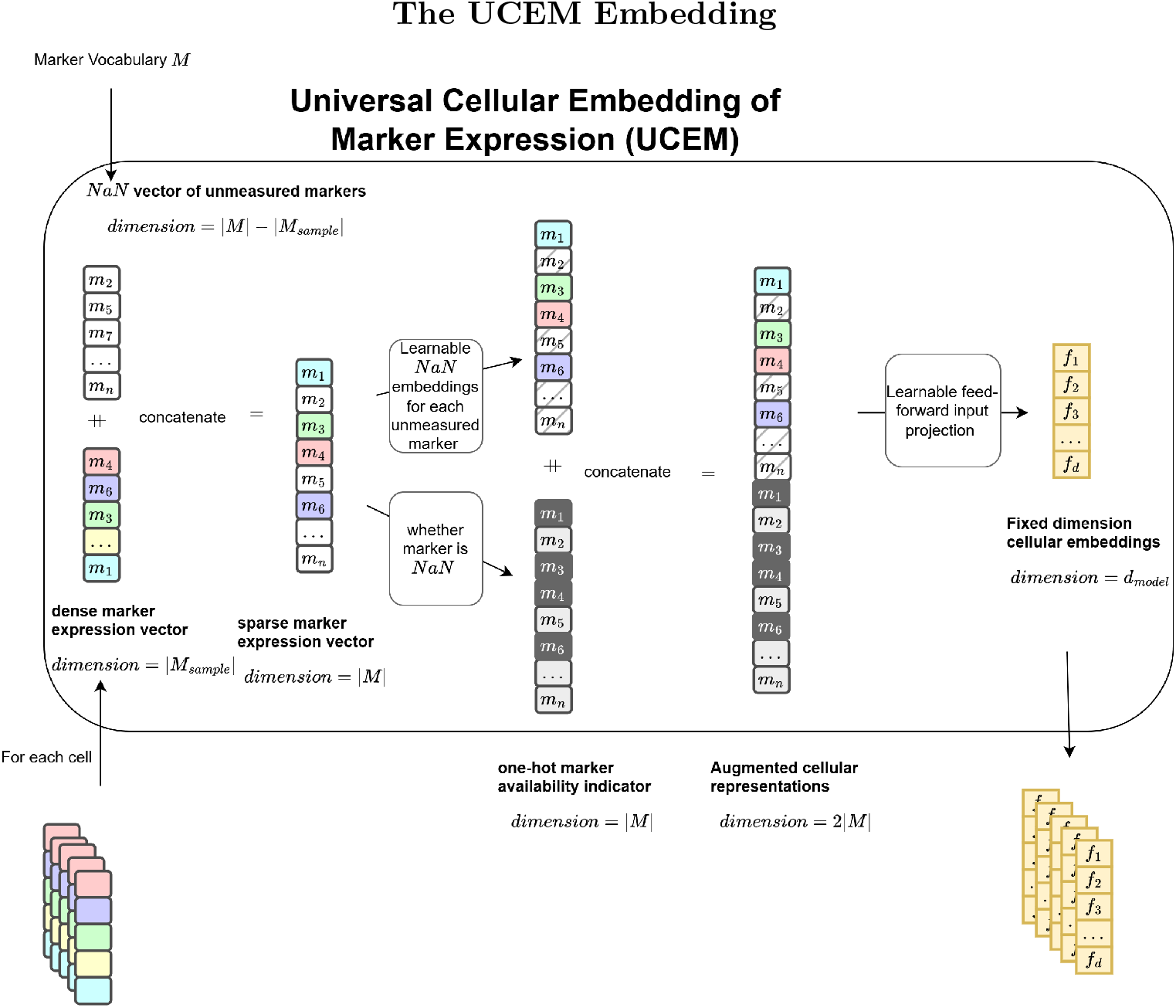
The UCEM embedding works by concatenating a sparse marker expression vector with a one-hot marker availability indicator to create an augmented representation of fixed dimension.

**Marker availability indicator** *a*_*i*_ ∈ {0, 1}^|*M* |^ as:

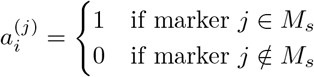

**Sparse marker expression vector** *e*_*i*_ ∈ ℝ^|*M* |^ as:

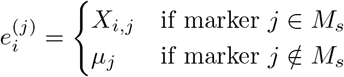

where *X*_*i,j*_ is the measured expression of *j* on cell *i*, and *µ*_*j*_ ∈ ℝ is a learnable masking value for unmeasured marker *j*.

#### Augmented cell representation

The augmented cell vector *c*_*i*_ is obtained through concatenation:

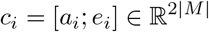

In the GPCT pipeline, *c*_*i*_ is then projected to the dimension specified by the transformer hidden size *h* through simple feed-forward layer(s).

This approach ensures that the model can distinguish unmeasured markers from measured but unexpressed markers, leveraging the learnable mask 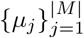 to capture biological priors for each marker type.

The UCEM embedding is conditionally robust to redundant markers not present in the training data, which always assume constant mask embedding values and are thus treated as redundant features, which the model learns to ignore.

#### 4.2.2 Transformer Encoder-Decoder

GPCT uses an encoder-decoder architecture (see in Supplementary Fig. 1), modified from the original Transformer proposed in [11]. The encoder-decoder transformer takes the set of embedded cell vectors generated by UCEM {*ci*}, and produces a sample-level prediction † as a output. No positional encoding was imposed due to lack of inherent order or spacial structures in observed cells, which is a common practice for data structures that require permutation invariance, such as the Set Transformer [16]. The encoder consists of a stack self attention blocks and operates solely on the tokens present in the sample. Each block contains a multi-headed scaled dot-product attention layer, followed by a two-layer position-wise feed-forward network. Layer normalisation is placed inside the residual connection as per Supplementary Fig. 1 in Pre-LN fashion [26] for better behaving gradients and ease of training.

Similarly, the decoder contains a stack of cross attention blocks which uses the cross attention between the contextualised encoder output (for Key and Value) and a task-specific learnable prediction token (for Query). The final state of the prediction token, being a task-conditioned representation of the sample, serves as the input to a feed-forward prediction head that predicts the sample-level label of the relevant downstream task, be it classification or regression.

The attention mechanism, in both cases, captures dynamic patterns from a set of tokens through learnable matrix multiplications that assign “attention” weights to each token, which allows the model to learn task-relevant patterns by dynamically weighting each token’s contribution, balancing global and local information in the process. The attention weights are often interpreted as token relevancy to learned patterns, giving transformer-based models inherent interpretability.

### 4.3 Training and optimisation

#### 4.3.1 Pretraining

In the pretraining stage, no external labels are used, with both the UCEM embedding and the GPCT-Encoder trained to optimise a suite of self-supervised cell-level masked prediction tasks. Three cell-level targets are derived from the raw data and used for self-supervision, including the per-marker expression values, as well as the percentile and the local density relative to sample-wide per-marker distribution. While it is known that self-supervision through reconstructing masked or corrupted inputs [27– 30] can extract robust and meaningful features, the percentile and density statistics are introduced specifically to tailor masked reconstruction towards flow cytometry data. Compared with the raw expression value, these two statistics reflect sample-wide distributions of marker expression on a per-cell basis, encouraging the model to learn sample-level cellular patterns despite cell-level self-supervision. Furthermore, the percentile and density statistics can be less sensitive to systematic signal shifts, directing the model’s focus towards patterns that are robust to experimental batch effects.

The pretraining stage is explained as follows, with a visualisation of the process available in Supplementary Fig 2.

##### Prediction targets

Given a sample’s marker expression matrix *X* ∈ ℝ^*n×m*^ where *n* is the number of cells and *m* = |*M*_*s*_| is the number of markers, we take *X* and compute two per-marker statistics *P* and *D* to obtain three prediction targets:

##### Percentile matrix *P*

For each marker *j* ∈ *{*1, …, *m}*, we compute the percentile rank of the expression for each cell:

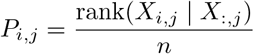

where rank(*X*_*i,j*_ |*X*_:,*j*_) denotes the rank of cell *i*’s expression for marker *j* among all *n* cells, yielding a percentile matrix *P* ∈ [0, 1]^*n×m*^.

##### Local density matrix *D*

For each marker *j*, we estimate the local density of each cell’s expression value based on the sample-wide distribution:

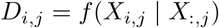

where *f* (· |*X*_:,*j*_) denotes the density function of marker *j*’s expression distribution, yielding a density matrix *D* ∈ ℝ^*n×m*^.

In this study, we implement *f* using logarithmically scaled kernel density estimation (KDE) with a Gaussian kernel:

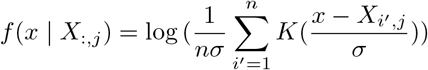

where *K* is the Gaussian density 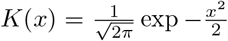 . The bandwidth hyperparameter *σ* is set to be 0.05 for all samples.

In practice, *P* and *D* are precomputed for each sample (up to 50,000 cells) for efficiency reasons.

#### Input masking

For each marker expression matrix *X* ∈ ℝ^*n×m*^, two complementary masking patterns are applied, yielding two masked expression matrices 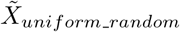 and 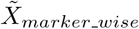 that both serve as input to the masked prediction task.

#### Uniform random masking

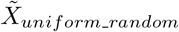 is created through uniform random masking per cell that follows an optional sample-wide marker dropout. First, for each cytometry sample, we randomly sample a dropout ratio *r*_*drop*_ ∼ Beta(*α*_1_, *β*_1_) and randomly remove expression values of *round*(*m* · *r*_*drop*_) markers by treating them as unmeasured across all cells. Then, a masking ratio *r*_*mask*_ ∼ Beta(*α*_2_, *β*_2_) is randomly sampled, and a masking matrix ℳ_*uniform*_*_*_*random*_is randomly generated as follows:

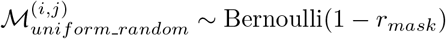

where 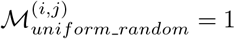 indicates that cell *i*’s expression for marker *j* is retained and 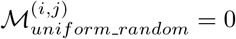 indicates masking. The resulting masked matrix is obtained as:

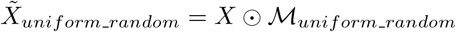

where ⊙ denotes element-wise multiplication. This yields a sparse matrix with uniformly distributed masked values, which encourages the model to explore inter-cell relationships for robust learned features.

##### Marker-wise masking

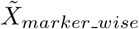 is created by masking expression values for selected markers across all cells. For each sample, a masking ratio *r*_*mask*_ ∼ Beta(*α*_3_, *β*_3_) is generated, and *round*(*m* · *r*_*mask*_) markers are selected as masked. A masking matrix ℳ_*marker*_*_*_*wise*_ is then generated as follows:

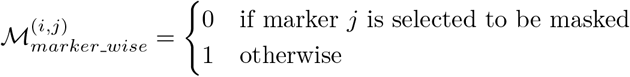

where 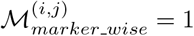 indicates that cell *i*’s expression for marker *j* is retained, and 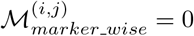 indicates masking. The resulting masked matrix is obtained as:

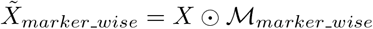

where ⊙ denotes element-wise multiplication. This yields a matrix where a marker is either masked for all cells or observed for all cells, mimicking the data distribution of an unmasked sample and encouraging the model to explore inter-marker correlations. The masked marker expressions are treated as unobserved during UCEM embedding, but distinguished from unobserved marker expressions when formulating the pretraining objective. Parameters for the beta distributions used in this study can be found in the supplementary information.

#### Pretraining Task Formulation

For each sample *X*, we apply both masking patterns, obtain the two pattern-specific objectives, before combining them to get the final pretraining objective.

For each masking pattern *k* ∈ {*uniform_random, marker_wise*}, the masked matrix 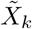 is passed through the UCEM embedding and GPCT-Encoder to obtain a latent representation *Z*_*k*_ at encoder output, from which three prediction heads jointly predict the targets:

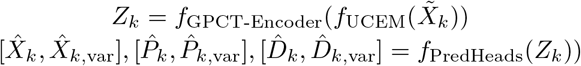

where 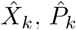, and 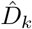 denote predicted marker expressions, percentiles, and densities respectively, and 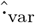 denotes predicted variance for each. Each prediction is computed over all markers in the vocabulary *M*, yielding tensors of shape *n* × |*M*| ; however, only measured markers contribute to the loss computation, as no ground truth exists for the unmeasured ones.

For each target *t* ∈ {*X, P, D* }, we compute the Gaussian negative log-likelihood (GNLL) loss separately on masked and unmasked entries, which are then summed up to produce the loss for the target:

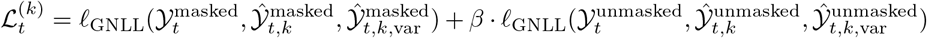

where the superscripts denote whether entries are masked or unmasked, and *β <* 1 is a regularisation weight that prevents the model from disregarding observed marker values.

The combined loss for masking pattern *k* across all three targets is:

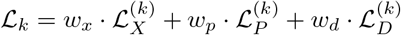

where *w*_*x*_, *w*_*p*_, and *w*_*d*_ are target weights. The final pretraining objective combines both masking patterns through a weighted sum:

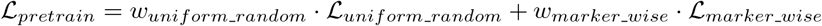

In this study, *w*_*x*_, *w*_*p*_, and *w*_*d*_ are all set to be 1, *w*_*uniform*_*_*_*random*_ and *w*_*marker*_*_*_*wise*_ are both set to be 0.5.

This dual-masking formulation drives the model to learn robust representations by predicting masked values from complementary perspectives: random masking encourages learning inter-cell relationships, while marker-wise masking promotes learning inter-marker correlations. The multi-task learning approach encourages the encoder to capture both local marker distributions and global cell-to-cell relationships, resulting in representations that are robust to measurement variability while preserving biological signals. Together, the pretraining stage guide the GPCT-Encoder towards projecting each cell from a noisy and heterogenous marker expression space into a consistent, biologically meaningful latent cell feature space.

#### 4.3.2 Downstream task-specific decoder training

Following pretraining, a GPCT-Decoder and the feed-forward task-specific prediction head is created to perform downstream predictions from the encoder output *Z* (Supplementary Fig. 1):

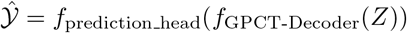

where 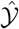 denotes the prediction, and the latent representation *Z* ∈ ℝ^*n×h*^ (*h* being the hidden dimension) is obtained from unmasked input *X* ∈ ℝ^*n×m*^:

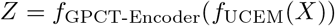

During downstream training, the parameters in the UCEM embedding and the GPCT-Encoder are frozen to preserve the learned cell representation, and a random marker-wise dropout is applied to the input sample *X* in place of encoder neuron dropout, derived in the same way as marker-wise masking in the pretraining stage, but with a fixed dropout rate, set to be 0.1 in this study. This procedure encourages the decoder to base its decisions on robust cellular patterns, avoid malignant overfitting to specific markers, and generalise better across different marker panels.

Similar to pretraining, the downstream training aims to reduce a task-specific loss:

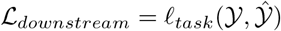

where 𝒴 denotes the ground truth labels, *ℓ*_*task*_ is the task-specific loss function (e.g., mean squared error (MSE) for regression or cross-entropy for classification), and ℒ_*downstream*_ is the computed loss value. In this study, cross-entropy is used as the loss function for classification.

In practice, we observed that in scenarios with abundant training data, unfreezing the UCEM embedding and the GPCT-Encoder to jointly optimise with the GPCT-Decoder could marginally improve downstream task performance. However, when data for downstream training is limited, freezing the encoder is recommended to maintain the generic cellular representations learned during pretraining.

### 4.4 Experimental settings

#### Training setups

The marker vocabulary used in this study contains 26 markers, which covers all markers in the ENU dataset and the KOMP dataset (full list in Supplementary Table 2).

All cross validation splits are stratified according to class label and grouped by experimental batches to avoid false confidence in model ability induced by unattended batch effects. For all biological sex prediction tasks, the training set is further divided into training and validation using a 90-10 split, with validation reserved for early stopping. For the gene KO prediction task, all models were trained to convergence on all of the training samples, due to the limited data quantity. The predictions are then aggregated from each fold and evaluated using accuracy, macro-averaged F1, and the Area Under Curve (AUC) score.

For all GPCT models, we used 4 encoder layers (when applicable) and 4 decoder layers, each with 4 attention heads. The hidden size of the model is set to be 128. Further model hyperparameters can be found in Supplementary Table 5. During training, we subsampled each cytometry sample down to 5000 cells for pretraining, and 7000 for downstream training. Further optimisation hyperparameters can also be found in Supplementary Table 4, 7, 6. CellCnn was run with default hyperparameters, unless otherwise listed in the supplementary information. All changes to the default hyperparameter values were to ensure convergence and fair comparison with GPCT.

#### Data preprocessing

During preprocessing of the FCS files, logicle transformation was applied to all samples. Channels with missing marker information were discarded, as were extremely rare markers appearing only once in the entire dataset. For the ENU dataset, 56 antibody-fluorophore conjugations were identified for 21 flow cytometry markers (see Supplementary Table 10), which makes up 16 marker panels (see Supplementary Table 8), most of which are largely pan-lymphoid. B220 and CD3 are the only two markers measured for all samples. For the KOMP dataset, all 10 markers were measured using consistent antibody-fluorophore conjugations (see Supplementary Table 3).

#### Model Training details

All models were implemented using PyTorch and trained using NVIDIA A100 or H200 GPUs. For reference, The mouse biological sex classification task using Datset 1 took 34.5 hours under 7-fold cross-validation (averaging 5 hours per two-stage training run) on an Nvidia H200 GPU with the specified hyperparameters, with an average pretraining epoch taking up around 50 seconds, and an average downstream training epoch taking up around 15 seconds. Each epoch contains approximately 170 steps. Inference speed was around 182 samples per second with each input sample containing 7000 cells, and is expected to scale quadratically with respect to the number of cells.

## 5 Code and Data

Dataset 2 was published by the data owner on FlowRepository, under the code FR-FCM-ZYPJ.

The de-identified Dataset 1 and weights of trained GPCT models can be found as supplementary data at https://doi.org/10.5281/zenodo.18975086

Source code for GPCT can be found on Github at https://github.com/ZoraZhuang/GPCT/

## 6 Acknowledgements

The authors thank the National Computational Infrastructure (Australia) for continued access to significant computation resources, and the Cytometry, Histology and Advanced Spatial Multiomics Facility at the John Curtin School of Medical Research. The authors gratefully acknowledge access to a substantial dataset of raw flow cytometry data from the mouse, generated by the Australian Phenomics Network at the Australian Phenomics Facility with financial support from the National Collaborative Research Infrastructure Strategy (Australia) and the National Institutes of Health (USA; Grant AI100627) to C. Goodnow. We additionally acknowledge A. Enders, L. Miosge, Y. Sontani and C. Roots in the production of this large, high-quality dataset.

## References

[1] Aghaeepour, N., Finak, G., Consortium, F., Consortium, D., Hoos, H., Mosmann, T.R., Brinkman, R., Gottardo, R., Scheuermann, R.H.: Critical assessment of automated flow cytometry data analysis techniques. Nature methods 10(3), 228–238 (2013)

[2] Hu, Z., Bhattacharya, S., Butte, A.J.: Application of machine learning for cytometry data. Frontiers in immunology 12, 787574 (2022)

[3] Arvaniti, E., Claassen, M.: Sensitive detection of rare disease-associated cell subsets via representation learning. Nature communications 8(1), 14825 (2017)

[4] Hu, Z., Tang, A., Singh, J., Bhattacharya, S., Butte, A.J.: A robust and interpretable end-to-end deep learning model for cytometry data. Proceedings of the National Academy of Sciences 117(35), 21373–21380 (2020)

[5] Weijler, L., Kowarsch, F., Reiter, M., Hermosilla, P., Maurer-Granofszky, M., Dworzak, M.: Fate: Feature-agnostic transformer-based encoder for learning generalized embedding spaces in flow cytometry data. In: Proceedings of the IEEE/CVF Winter Conference on Applications of Computer Vision, pp. 7956–7964 (2024)

[6] Wödlinger, M., Reiter, M., Weijler, L., Maurer-Granofszky, M., Schumich, A., Sajaroff, E.O., Groeneveld-Krentz, S., Rossi, J.G., Karawajew, L., Ratei, R., et al.: Automated identification of cell populations in flow cytometry data with transformers. Computers in Biology and Medicine 144, 105314 (2022)

[7] Lewis, J.E., Cooper, L.A., Jaye, D.L., Pozdnyakova, O.: Automated deep learning-based diagnosis and molecular characterization of acute myeloid leukemia using flow cytometry. Modern Pathology 37(1), 100373 (2024)

[8] Robles, E.E., Jin, Y., Smyth, P., Scheuermann, R.H., Bui, J.D., Wang, H.-Y., Oak, J., Qian, Y.: A cell-level discriminative neural network model for diagnosis of blood cancers. Bioinformatics 39(10), 585 (2023)

[9] Yi, H., Stanley, N.: Cytoset: Predicting clinical outcomes via set-modeling of cytometry data. In: Proceedings of the 12th ACM International Conference on Bioinformatics, Computational Biology, and Health Informatics, pp. 1–8 (2021)

[10] Cox, A.M., Kim, D., García, R., Fuda, F.S., Weinberg, O.K., Chen, W.: Automated prediction of acute promyelocytic leukemia from flow cytometry data using a graph neural network pipeline. American journal of clinical pathology 161(3), 264–272 (2024)

[11] Vaswani, A., Shazeer, N., Parmar, N., Uszkoreit, J., Jones, L., Gomez, A.N., Kaiser, Ł., Polosukhin, I.: Attention is all you need. Advances in neural information processing systems 30 (2017)

[12] Minaee, S., Mikolov, T., Nikzad, N., Chenaghlu, M., Socher, R., Amatriain, X., Gao, J.: Large language models: A survey. arXiv preprint 2402.06196 (2024)

[13] Szalata, A., Hrovatin, K., Becker, S., Tejada-Lapuerta, A., Cui, H., Wang, B., Theis, F.J.: Transformers in single-cell omics: a review and new perspectives. Nature methods 21(8), 1430–1443 (2024)

[14] Abramson, J., Adler, J., Dunger, J., Evans, R., Green, T., Pritzel, A., Ronneberger, O., Willmore, L., Ballard, A.J., Bambrick, J., et al.: Accurate structure prediction of biomolecular interactions with alphafold 3. Nature 630(8016), 493–500 (2024)

[15] Zhao, W.X., Zhou, K., Li, J., Tang, T., Wang, X., Hou, Y., Min, Y., Zhang, B., Zhang, J., Dong, Z., et al.: A survey of large language models. arXiv preprint 2303.18223 (2023)

[16] Lee, J., Lee, Y., Kim, J., Kosiorek, A., Choi, S., Teh, Y.W.: Set transformer: A framework for attention-based permutation-invariant neural networks. In: International Conference on Machine Learning, pp. 3744–3753 (2019). PMLR

[17] Kowarsch, F., Weijler, L., Wödlinger, M., Reiter, M., Maurer-Granofszky, M., Schumich, A., Sajaroff, E.O., Groeneveld-Krentz, S., Rossi, J.G., Karawajew, L., et al.: Towards self-explainable transformers for cell classification in flow cytometry data. In: International Workshop on Interpretability of Machine Intelligence in Medical Image Computing, pp. 22–32 (2022). Springer

[18] Nelms, K.A., Goodnow, C.C.: Genome-wide enu mutagenesis to reveal immune regulators. Immunity 15(3), 409–418 (2001)

[19] Mashford, B.S., Hewitt, T., May, M., Chuah, A., Andrews, D.: Comparison of deep-learning models for classification of cellular phenotype from flow cytometry data. IEEE Transactions on Computational Biology and Bioinformatics 22(4), 1587–1592 (2025) 10.1109/TCBBIO.2025.3562597

[20] Bradley, A., Anastassiadis, K., Ayadi, A., Battey, J.F., Bell, C., Birling, M.-C., Bottomley, J., Brown, S.D., Bürger, A., Bult, C.J., et al.: The mammalian gene function resource: the international knockout mouse consortium. Mammalian genome 23(9), 580–586 (2012)

[21] Abeler-Dörner, L., Laing, A.G., Lorenc, A., Ushakov, D.S., Clare, S., Speak, A.O., Duque-Correa, M.A., White, J.K., Ramirez-Solis, R., Saran, N., et al.: High-throughput phenotyping reveals expansive genetic and structural underpinnings of immune variation. Nature Immunology 21(1), 86–100 (2020)

[22] McInnes, L., Healy, J., Melville, J.: Umap: Uniform manifold approximation and projection for dimension reduction. arXiv preprint 1802.03426 (2018)

[23] Mashford, B.S., Hewitt, T., May, M., Zhuang, Z., Jain, A., Diamand, K.E., Li, F.-J., Kwong, K., Read, S.H., Davies, A.R., et al.: Superior batch alignment and hyper-dimensional cytometry representations allow ultra-sensitive classification of disease phenotypes. bioRxiv, 2025–07 (2025)

[24] Chefer, H., Gur, S., Wolf, L.: Transformer interpretability beyond attention visualization. In: Proceedings of the IEEE/CVF Conference on Computer Vision and Pattern Recognition, pp. 782–791 (2021)

[25] Chefer, H., Gur, S., Wolf, L.: Generic attention-model explainability for interpreting bi-modal and encoder-decoder transformers. In: Proceedings of the IEEE/CVF International Conference on Computer Vision, pp. 397–406 (2021)

[26] Xiong, R., Yang, Y., He, D., Zheng, K., Zheng, S., Xing, C., Zhang, H., Lan, Y., Wang, L., Liu, T.: On layer normalization in the transformer architecture. In: International Conference on Machine Learning, pp. 10524–10533 (2020). PMLR

[27] Vincent, P., Larochelle, H., Bengio, Y., Manzagol, P.-A.: Extracting and composing robust features with denoising autoencoders. In: Proceedings of the 25th International Conference on Machine Learning, pp. 1096–1103 (2008)

[28] Devlin, J., Chang, M.-W., Lee, K., Toutanova, K.: Bert: Pre-training of deep bidirectional transformers for language understanding. In: Proceedings of the 2019 Conference of the North American Chapter of the Association for Computational Linguistics: Human Language Technologies, Volume 1 (long and Short Papers), pp. 4171–4186 (2019)

[29] Bao, H., Dong, L., Piao, S., Wei, F.: Beit: Bert pre-training of image transformers. arXiv preprint 2106.08254 (2021)

[30] Zhang, C., Zhang, C., Song, J., Yi, J.S.K., Zhang, K., Kweon, I.S.: A survey on masked autoencoder for self-supervised learning in vision and beyond. arXiv preprint 2208.00173 (2022)

