## Supplementary Information for "Towards a Cytometry Foundation Model: Interpretable Sample-level Predictive Modelling via Pretrained Transformers"

### 1 Further information on modelling mouse biological sex across Dataset 1 and 2

Dataset 2 is split into a small training set containing 124 samples taken from 4 experimental batches (Dataset 2 training data) , and a large testing set containing the rest of labelled samples, totalling 6668 (Dataset 2 testing data). The Dataset 2 testing data is reserved for testing for all experiments, and the Dataset 2 training data may be provided in unlabelled or labelled format, depending on the setup.

To ensure optimal performance during generalisation, markers specific to the target dataset (Dataset 2) should be excluded if they were not present in the source training set (Experiment B). As these markers were consistently masked during all training phases, the model could not derive useful representations for them. Attempting to utilise their unmasked values during inference (Experiment C) introduces out-of-distribution noise, which significantly degrades performance, often falling below a random baseline, as shown in Figure 2b (1) in the main text.

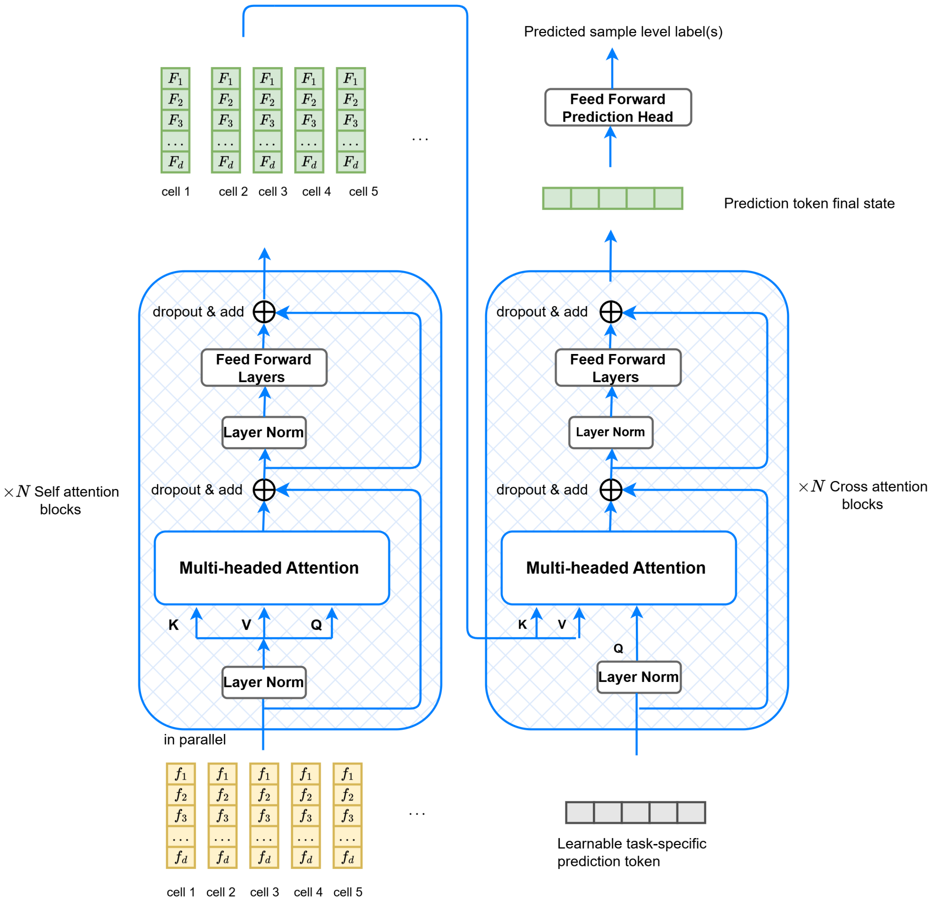


**Fig. 1** The Encoder-Decoder architecture of GPCT. No positional encoding is imposed, and layer norms are placed inside the residual connections (Pre-LN style). A learnable task-specific prediction token is utilised by the decoder to produce a sample level prediction.

**Table 1** Performance of each model for the gene KO prediction task. Metrics are calculated based on aggregated leave-one-batch-out cross validation results. AUC (ovo) and AUC (ovr) stand for macro-average one-versus-one and macro-average one-versus-rest AUC scores, respectively.

| Model | ACC | Macro-average F1 | AUC (ovo) | AUC (ovr) |
|---|---|---|---|---|
| GPCT | **0.694** | **0.683** | **0.903** | **0.901** |
| GPCT decoder only | 0.431 | 0.427 | 0.735 | 0.733 |
| CellCnn | 0.458 | 0.450 | 0.725 | 0.726 |
| GPCT generic encoder | ***0.736*** | ***0.737*** | ***0.921*** | ***0.919*** |

**Table 2** Marker vocabulary used in this study. Markers in italics are observed in the ENU dataset, bold markers are observed in the KOMP dataset. Bold italics are observed in both.

| **Marker** | |
|---|---|
| *ab-TCR* | **CD45-2** |
| **B220** | *CD62L* |
| **CD1d** | ***CD8*** |
| **CD21** | **Foxp3** |
| **CD23** | *gd-TCR* |
| ***CD25*** | **IgD** |
| **CD3** | **IgE** |
| **CD365** | **IgM** |
| ***CD4*** | ***KLRG1*** |
| ***CD44*** | **NK1-1** |
| *CD45* | **CD127** |
| **CD93** | **CD122** |
| **CD138** | *CD161* |

**Table 3** KOMP marker-fluorophore combinations.

| **Fluorophore** | **Marker** |
|---|---|
| APC-A | gd-TCR |
| APC-Cy7-A | CD8 |
| Alexa Fluor 700-A | CD45 |
| AmCyan-A | CD4 |
| FITC-A | CD44 |
| PE-A | CD25 |
| PE-Cy7-A | KLRG1 |
| PE-Texas Red-A | CD62L |
| Pacific Blue-A | CD161 |
| PerCP-Cy5-5-A | ab-TCR |

**Table 4** Shared hyperparameters for GPCT model optimisation, unless otherwise specified. Decoder only variants have reduced dropout ratio to reduce possibility of failure to learn at initialisation.

| hyperparameter | default value |
|---|---|
| batch size | 64 |
| optimiser | AdamW |
| weight decay pretraining | 0.005 |
| weight decay downstream | 0.001 |
| pretraining learning rate | 5e-4 |
| pretraining epochs | 300 |
| pretraining cell tokens per sample | 5000 |
| pretraining Beta($\alpha_1, \beta_1$) | Beta(1.5,9) |
| pretraining Beta($\alpha_2, \beta_2$) | Beta(5,16) |
| pretraining Beta($\alpha_3, \beta_3$) | Beta(5,15) |
| pretraining $\beta$* | 0.1 |
| downstream learning rate | 2e-4 |
| downstream epochs | 200 (patience 50) |
| downstream cell tokens per sample | 7000 |
| dropout | 0.2 (0.1 for decoder only variants) |

**Table 5** Shared hyperparameters of model architecture for all GPCT models.

| hyperparameter | default value |
|---|---|
| marker vocabulary size | 26 |
| hidden layer size | 128 |
| encoder layers (if applicable) | 4 |
| decoder layers | 4 |
| attention heads | 4 |
| input projection | [linear(128), relu ] |
| classification head | [256,128,$n\_classes$] |
| all prediction heads in pretraining | [128 (shared by all heads), 52, 26] |

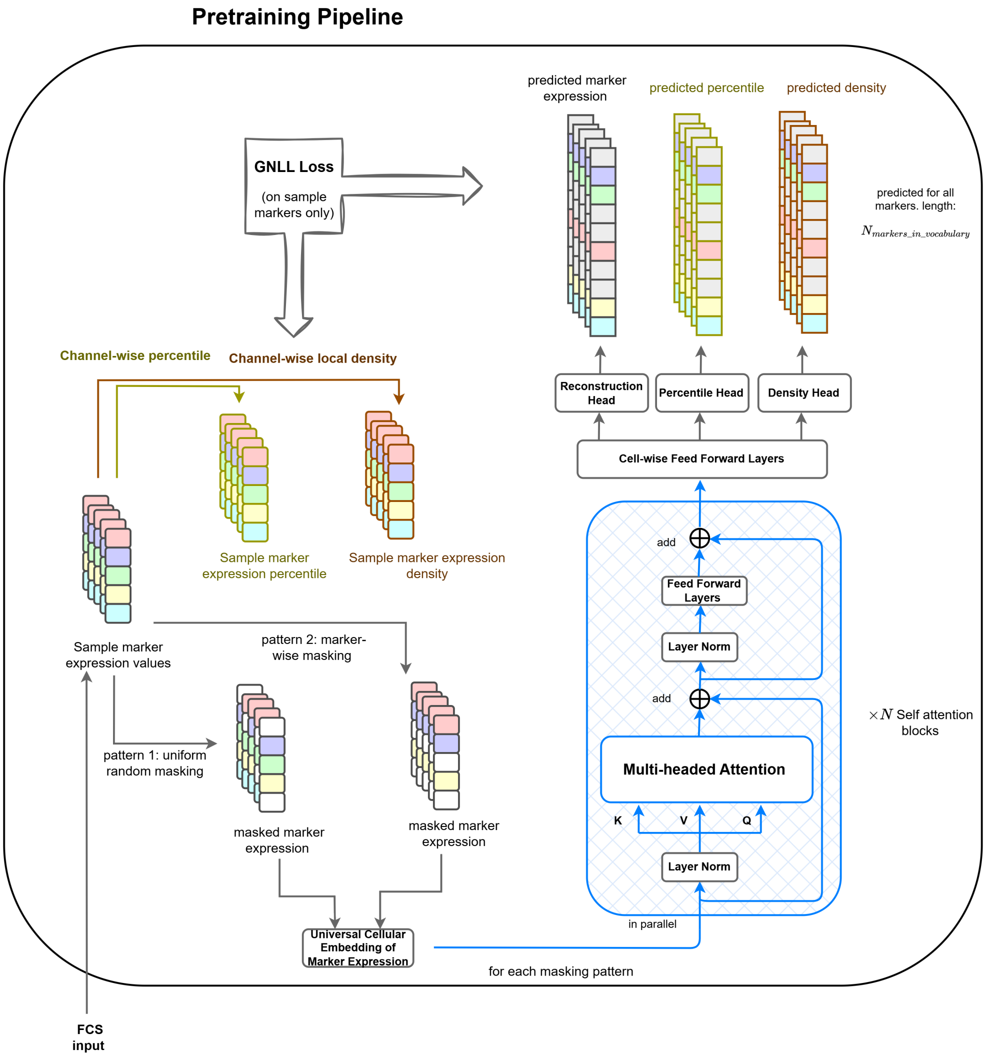


**Fig. 2** The self-supervised pretraining pipeline designed for robust cell representation learning: (1) the Percentile and Density matrices are calculated based on raw marker expression (2) The marker expression matrix is masked using two masking patterns, resulting in two masked marker expression matrices. For each masked matrix: (3) The mean and variance predictions of unmasked marker expression, percentile and density matrices for the whole marker vocabulary are made through the [UCEM embedding, GPCT-Encoder, cell-wise prediction heads] pipeline (4) the GNLL loss is calculated for both masked and unmasked entries, but not for initially unmeasured markers (no ground truth available) (5) the loss for the two masking patterns, each with three prediction tasks, are combined using pre-specified weights. Backpropagation is performed on the final loss, and the model parameters are updated.

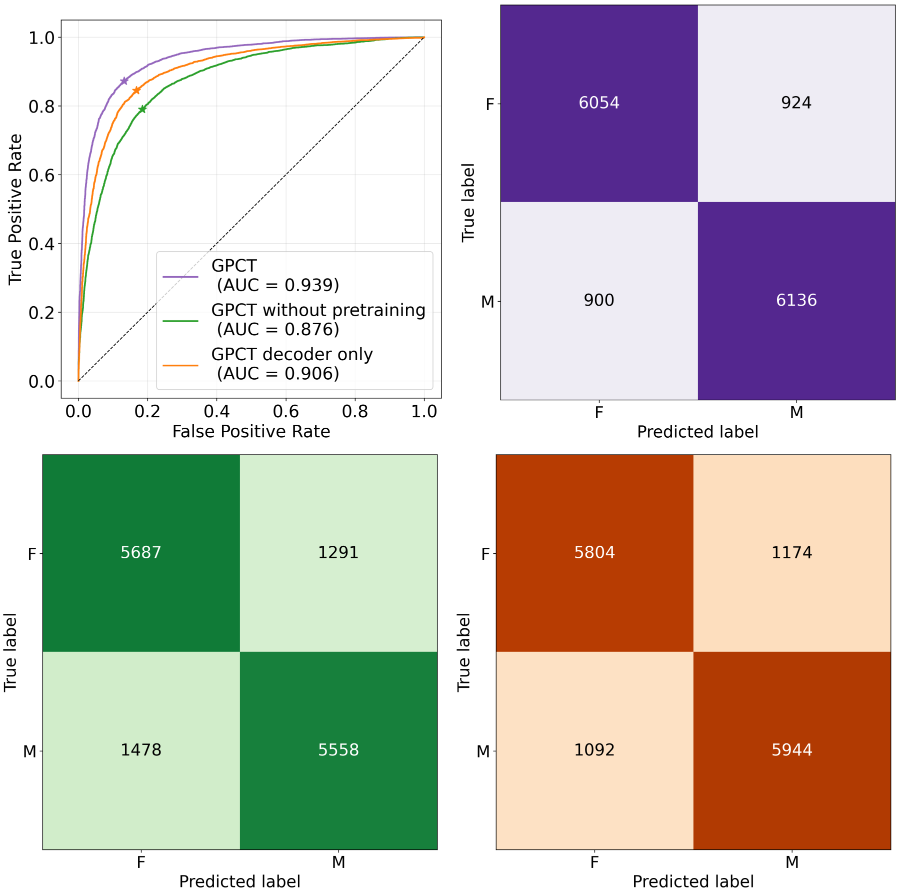


**Fig. 3** Results for the ENU mouse biological sex classification: Top left: ROC curves of GPCT, GPCT without pretraining, GPCT decoder only. Top right, bottom left, bottom right: Confusion matrices for GPCT, GPCT without pretraining, and GPCT decoder only. Generated based on aggregated 7-fold cross validation results.

**Table 6** Modelling biological sex across KOMP and ENU: Optimisation hyperparameters varied from default values for KOMP-data-only training (KOMP encoder and/or KOMP decoder). KOMP training samples were also upsampled 20 fold when training alongside ENU samples to account for differences in sample support.

| hyperparameter | value for KOMP only training |
|---|---|
| batch size | 16 |
| pretraining epochs | 2000 |
| downstream epochs | 400 |

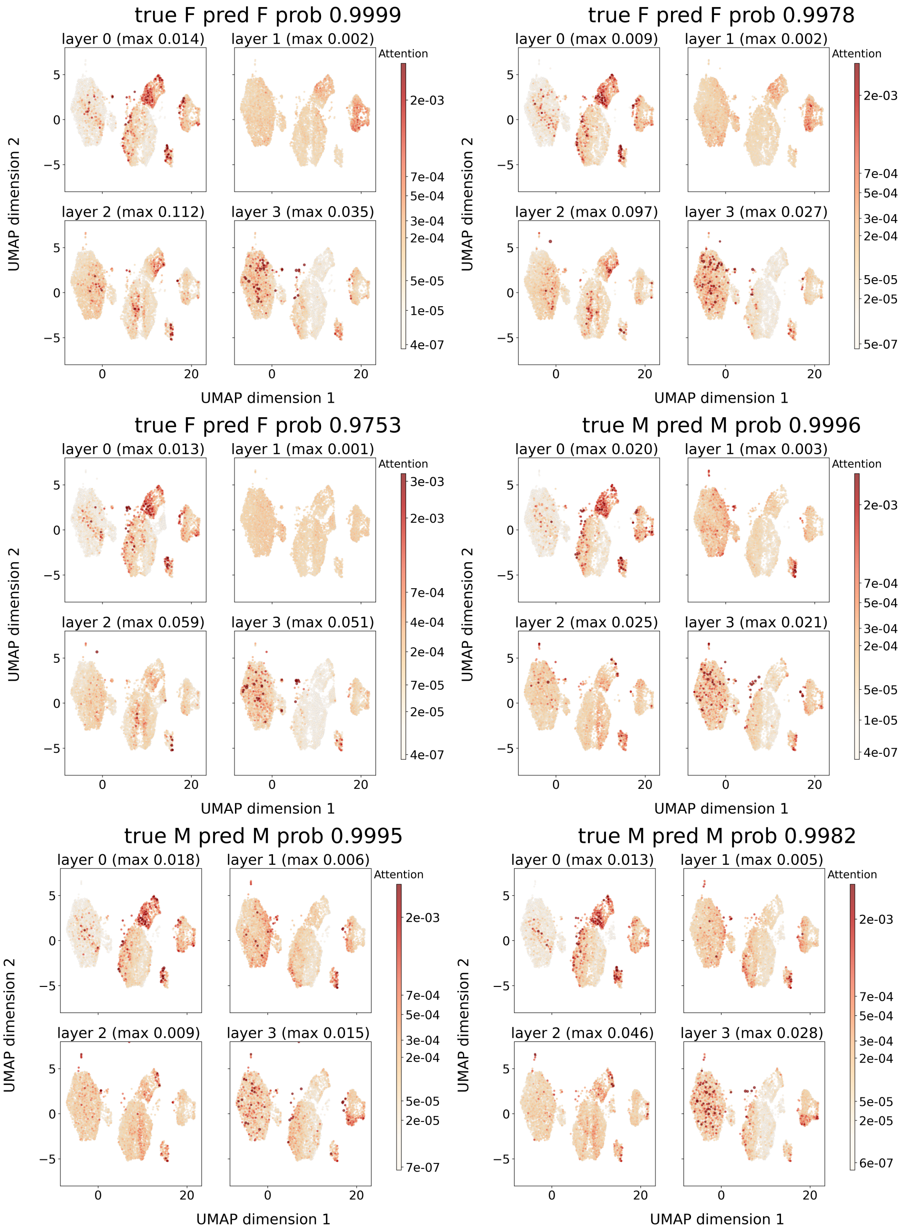


**Fig. 4** UMAP visualisation of attention per layer of 6 different ENU mice for the biological sex classification task. The title for each describes the true sex, the predicted sex, and the probability / confidence of the prediction. In general, attention remains consistent across samples of the same sex, and differs slightly for female and male samples, most notably in layer 1.

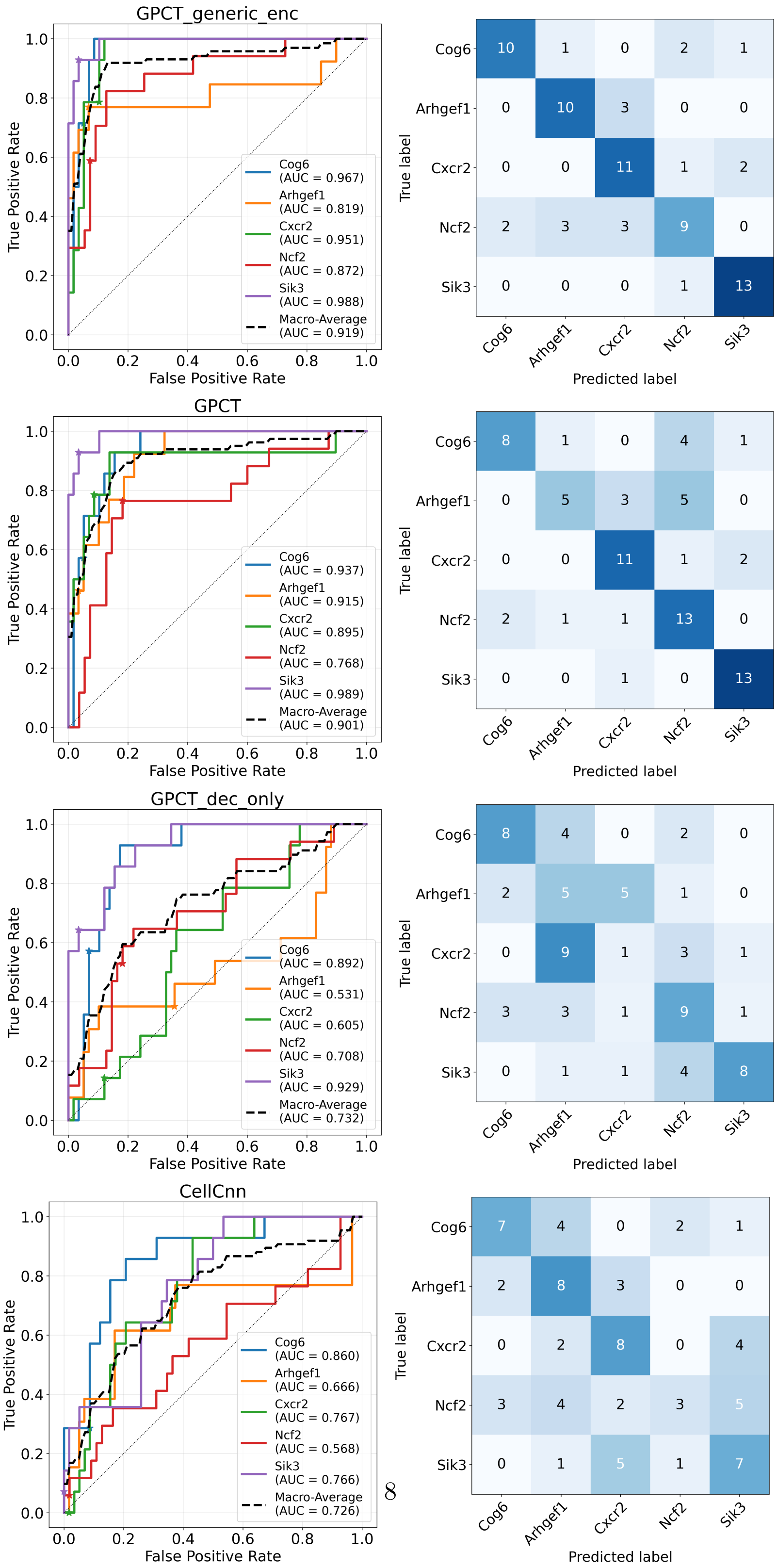


**Fig. 5** KOMP gene knockout prediction: The per-class and macro-averaged one-vs-rest ROC curves, as well as the confusion matrix for each of the four models (GPCT generic encoder, GPCT, GPCT decoder only, CellCnn). The star marker denotes the classification threshold of 0.5.

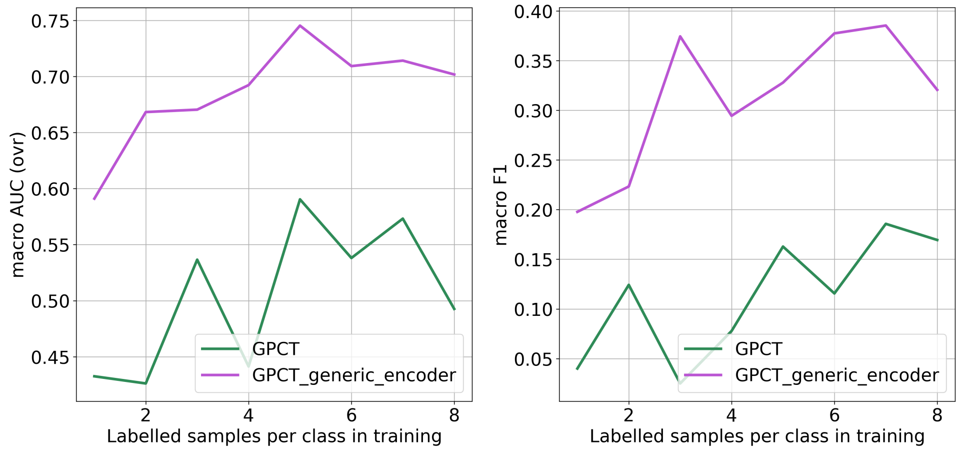


(a) (b)

**Fig. 6** Few-shot KO classification experiment on the KOMP dataset, measured by (a) macro-average one-vs-rest AUC and (b) macro-average F1 score. The dataset is split into train ($8 \times 5 = 40$ samples) and test (32) sets, with samples from the same batch kept within the same split to prevent shortcut learning from batch effects. All downstream training ran for 3000 steps at a batch size of 5. Pretraining ran for 6000 steps at a batch size of 5 for GPCT with non-generic encoder, while GPCT with generic encoder used the same generic encoder from the leave-one-batch-out cross validation.

**Table 7** Optimisation hyperparameters for KOMP KO predictions that are different from the default, mainly to account for a smaller number of training data. Hyperparameters marked with * do not apply to CellCnn.

| hyperparameter | value |
|---|---|
| batch size | 8 (32 for generic encoder) |
| pretraining epochs | 1000 (200 for generic encoder) |
| weight decay downstream* | 0.01 |
| downstream epochs | 200 (1000 for decoder only), patience 100 |
| pretraining $\beta$* | 0.05 |

**Table 8** Distinct marker panels in the ENU dataset and their sample counts

| **Marker Panel** | **Count** |
|---|---|
| B220,CD3,CD4,CD44,IgD,IgM,KLRG1,NK1-1 | 7002 |
| B220,CD25,CD3,CD4,CD44,IgD,IgM,NK1-1 | 6013 |
| B220,CD3,CD4,CD44,IgD,IgE,IgM,KLRG1,NK1-1 | 395 |
| B220,CD3,CD4,CD44,IgD,IgE,IgM | 178 |
| B220,CD21,CD23,CD3,CD93,IgD,IgM | 122 |
| B220,CD1d,CD3,CD4,CD44,NK1-1 | 65 |
| B220,CD127,CD3,CD4,CD44,CD8,IgD,IgM | 45 |
| B220,CD122,CD127,CD3,CD4,CD44,CD8,KLRG1 | 44 |
| B220,CD127,CD3,CD4,CD44,CD8 | 41 |
| B220,CD138,CD3,CD93,IgD,IgM | 31 |
| B220,CD138,CD21,CD23,CD3,CD365,CD93,IgD,IgM | 27 |
| B220,CD25,CD3,CD4,CD44,CD8,Foxp3 | 25 |
| B220,CD3,CD4,CD44,CD45-2,IgD,IgM,KLRG1,NK1-1 | 23 |
| B220,CD25,CD3,CD4,CD44,IgD,IgM | 1 |
| B220,CD25,CD3,CD44,IgD,IgM,NK1-1 | 1 |
| B220,CD3,CD4,CD44,CD45-2,CD8 | 1 |
| Total | 14014 |

**Table 9** Sample support and zygosity for each gene KO class.

| Gene | Cog6 | Arhgef1 | Cxcr2 | Ncf2 | Sik3 | Total |
|---|---|---|---|---|---|---|
| *n_samples* | 14 | 13 | 14 | 17 | 14 | 72 |
| Zygosity | Hom | Hom | Het | Hom | Hom | |

**Table 10**: ENU Marker-dye combination counts with marker totals

| Marker — Dye Combination | Count | Total for Marker |
|---|---|---|
| B220 — APC-A | 64 | 14014 |
| B220 — APC-Alexa 750 / APC-Cy7-A | 149 | |
| B220 — FITC-A | 25 | |
| B220 — PerCP-Cy5-5-A | 13776 | |
| CD122 — FITC-A | 44 | 44 |
| CD127 — PE-A | 110 | 130 |
| CD127 — PerCP-Cy5-5-A | 20 | |
| CD138 — PE-A | 27 | 58 |
| CD138 — Qdot 605-A | 31 | |
| CD1d — APC-A | 65 | 65 |
| CD21 — PerCP-Cy5-5-A | 149 | 149 |
| CD23 — Alexa 405 / Pac Blue-A | 149 | 149 |
| CD25 — APC-A | 25 | 6040 |
| CD25 — PE-Cy7-A | 6015 | |
| CD3 — Alexa Fluor 700-A | 197 | 14014 |
| CD3 — APC-Alexa 750 / APC-Cy7-A | 13634 | |
| CD3 — APC-Cy7-A | 142 | |
| CD3 — FITC-A | 41 | |
| CD365 — Qdot 605-A | 27 | 27 |
| CD4 — Alexa Fluor 700-A | 13795 | 13833 |
| CD4 — APC-Alexa 750 / APC-Cy7-A | 17 | |
| CD4 — PE-A | 21 | |
| CD44 — Alexa 405 / Pac Blue-A | 13489 | 13834 |
| CD44 — Alexa Fluor 405-A | 71 | |
| CD44 — APC-A | 178 | |
| CD44 — Pacific Blue-A | 71 | |
| CD44 — Qdot 605-A | 25 | |
| CD45-2 — APC-A | 1 | 24 |
| CD45-2 — Qdot 605-A | 23 | |
| CD8 — APC-A | 66 | 156 |
| CD8 — APC-Alexa 750 / APC-Cy7-A | 20 | |
| CD8 — PE-Cy7-A | 26 | |
| CD8 — PerCP-Cy5-5-A | 44 | |
| CD93 — APC-A | 180 | 180 |

Continued on next page...

Table 10 – continued

| **Marker — Dye Combination** | **Count** | **Total for Marker** |
|---|---|---|
| Foxp3 — Alexa 405 / Pac Blue-A | 25 | 25 |
| IgD — FITC-A | 225 | 13838 |
| IgD — PE-A | 13613 | |
| IgE — Qdot 605-A | 573 | 573 |
| IgM — FITC-A | 13613 | 13838 |
| IgM — PE-A | 31 | |
| IgM — PE-Cy7-A | 194 | |
| KLRG1 — PE-Cy7-A | 7464 | 7464 |
| NK1-1 — APC-A | 13434 | 13499 |
| NK1-1 — FITC-A | 44 | |
| NK1-1 — PE-Cy7-A | 21 | |
